# Clinically-weighted transcriptomic signatures for protein kinase inhibitor associated cardiotoxicity

**DOI:** 10.1101/075754

**Authors:** JGC van Hasselt, J Hansen, Y Xiong, J Shim, A Pickard, G Jayaraman, B Hu, M Mahajan, J Gallo, EA Sobie, MR Birtwistle, EU Azeloglu, R Iyengar

**Affiliations:** Department of Pharmacological Sciences and Systems Biology Center New York, Icahn School of Medicine at Mount Sinai, New York, NY, United States.; Department of Pharmacology, Leiden University, Leiden, Netherlands.; Department of Genetics and Genomic Sciences, Icahn School of Medicine at Mount Sinai, New York, NY, United States.

## Abstract

Cardiotoxicity (CT) involving diminished cardiac contractility and heart failure is a major adverse event associated with otherwise efficacious protein kinase inhibitors (KIs). Here, we sought to develop clinically-weighted transcriptomic signatures to predict risk of CT and to better understand the biological processes associated with CT risk. We obtained transcriptome-wide response profiles in four human primary cardiomyocyte cell lines that were treated with 22 different KIs using mRNA sequencing with 3’ digital gene expression. The FDA Adverse Event Reporting System was used to derive relative risk scores for four types of CT for different KIs. We used elastic net regression to associate these transcriptomic profiles with KI-associated risk scores for CT subtypes to obtain clinically-weighted transcriptomic signatures, which showed good predictive properties (cross-validation R^2^ >0.87). Our clinically-weighted transcriptomic signatures for KI-associated CT may be of relevance in early drug development for the prediction of KI-associated CT.

## INTRODUCTION

Protein kinase inhibitors (KIs) are a clinically important class of anticancer agents^1,2^. There are currently more than 28 KIs^3^ on the market, and more than 100 KIs in clinical development^4^. There are significant concerns regarding the safety profile of KIs. Cardiotoxicity (CT) is one clinically important adverse event associated with some KIs^5–7^. Typically, KI-associated CT manifests as loss of contractile function, which can subsequently lead to heart failure^8^. The human ‘kinome’ consists of more than 500 protein kinases^9^. Given that many KIs are not highly selective in the protein kinases they target^10^, the inhibition of any of these kinases in healthy cell types, including cardiomyocytes, may potentially lead to adverse drug effects such as CT^11^. For specific KIs, *in vitro* and *in vivo* studies have identified potential mechanisms for CT^12^, including mitochondrial function ^6,13,14^, endoplasmic reticulum stress response^14^ and AMPK inhibition^15^. Overall, however, the mechanisms of KI-induced CT are still poorly understood^12^. Furthermore, characterization of the link between clinically observed CT phenotypes and transcriptomic changes in cardiomyocytes could have tremendous prognostic value. Therefore, there is a need to better characterize the molecular changes of KI associated CT on a genome-wide basis.

The use of a systems pharmacology approach to understand and predict adverse events has received increasing interest^16,17^, and several successful case examples have been described^18,19^. Such approaches typically rely on appropriate global characterization of drug-induced molecular changes in affected cells and tissues of interest. However, for KI-associated CT, a systematic assessment of molecular changes associated with KI treatment is lacking. Addressing this gap, as part of the NIH-funded Library of Integrated Network Based Cellular Signatures (LINCS), the Drug Toxicity Signature Generation Center (DToxS; www.dtoxs.org) is currently generating large-scale transcriptomics and proteomics profiling datasets in clinically-relevant, human-derived cell lines, which are perturbed with large numbers of different drugs, including cardiomyocytes and KIs. Here, we describe how transcriptomic profiles of 22 KIs in four human primary cardiomyocyte cell lines were analyzed in conjunction with clinical data from the FDA Adverse Event Reporting System, in order to derive clinically-weighted transcriptomic signatures that may allow ranking prospective KIs for their relative risk of CT.

## RESULTS

### Limited overlap of differentially expressed genes across different KIs in cardiomyocytes

We used mRNA sequencing with 3’-digital gene expression generated from four primary human cardiomyocyte cell lines that were treated with up to 22 KIs at their respective maximum therapeutic concentrations for 48 h (Table 1, Table S3) for our analyses. Mean differential gene expression fold-change values were computed across cell lines and replicates for all KIs. After ranking by absolute gene expression fold-change and keeping the top 500 genes, we computed the Jaccard index across these KI-associated top ranked gene expression profiles, which indicated limited similarity given the Jaccard index of <0.25 (Figure 1A). The genes that were most commonly present in the top 500 ranked gene list across drugs ranked by their frequency of occurrence is shown in Figure 1B, i.e. gene TC2N was present in these top ranking genes for all KIs except for cabozantinib. Principal component analysis showed variable gene expression patterns for 9 KIs, while for the remaining KIs little variation in gene expression was seen (Figure 1C), where PC3 appeared most discriminating.

**Figure 1A:**
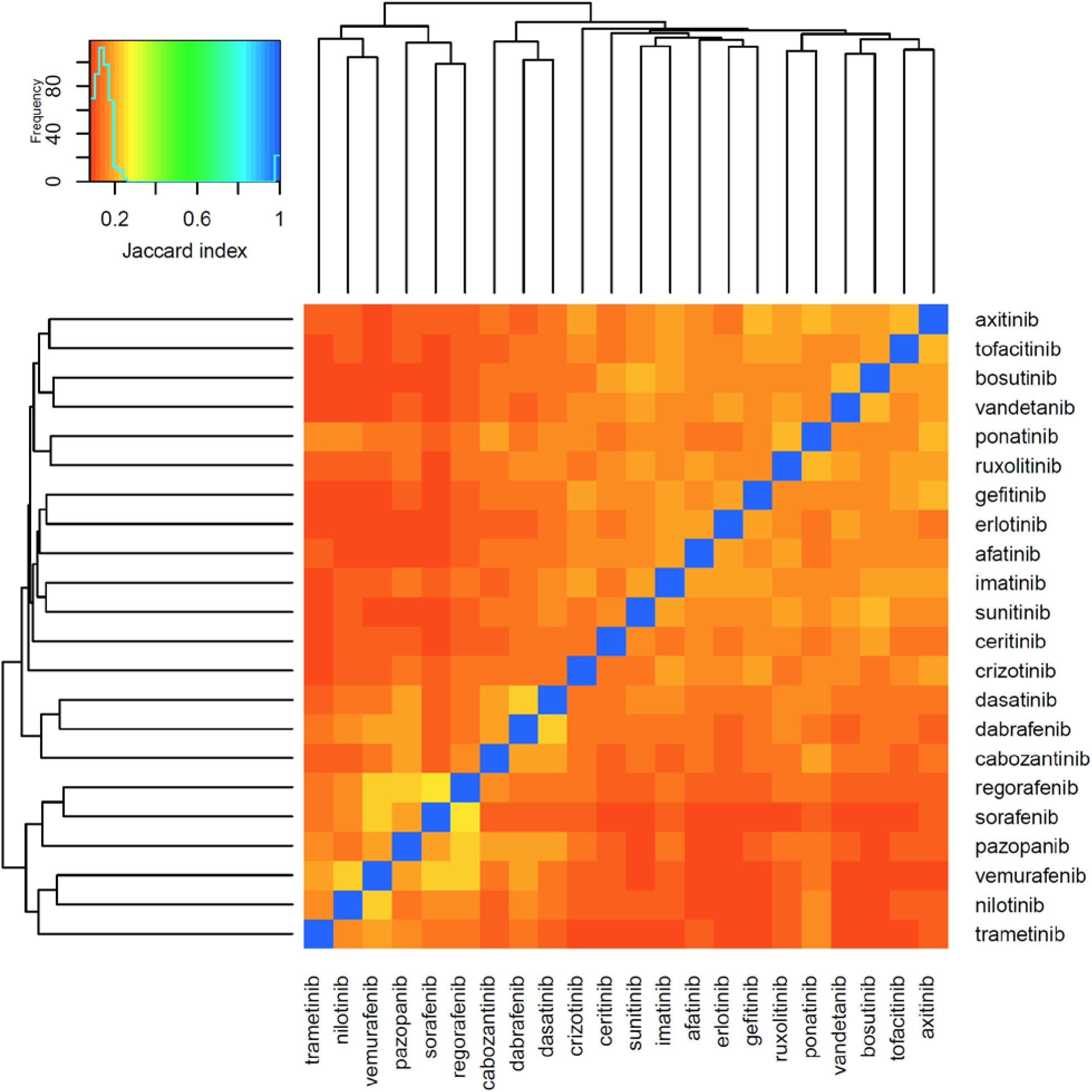
Transcriptomic profiling across KIs. For each drug, genes were ranked by mean fold-change gene expression value across replicates (>3) and cell lines, and the top 500 genes were kept. (A) Heatmap showing the Jaccard index which indicates the magnitude of similarity in top-ranking differentially expressed genes for all KI pairs.

**Figure 1B:**
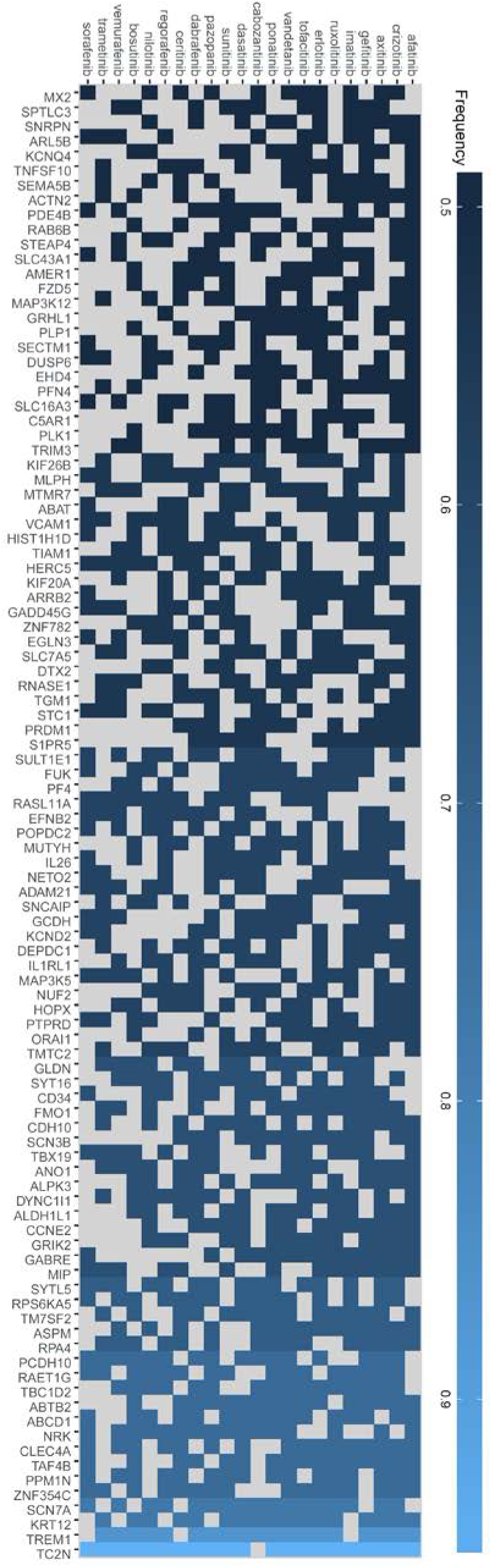
Transcriptomic profiling across KIs. For each drug, genes were ranked by mean fold-change gene expression value across replicates (>3) and cell lines, and the top 500 genes were kept. (B) Heatmap showing frequency of genes present for different KIs, with the genes and drugs ranked by frequency of occurrence

**Figure 1C:**
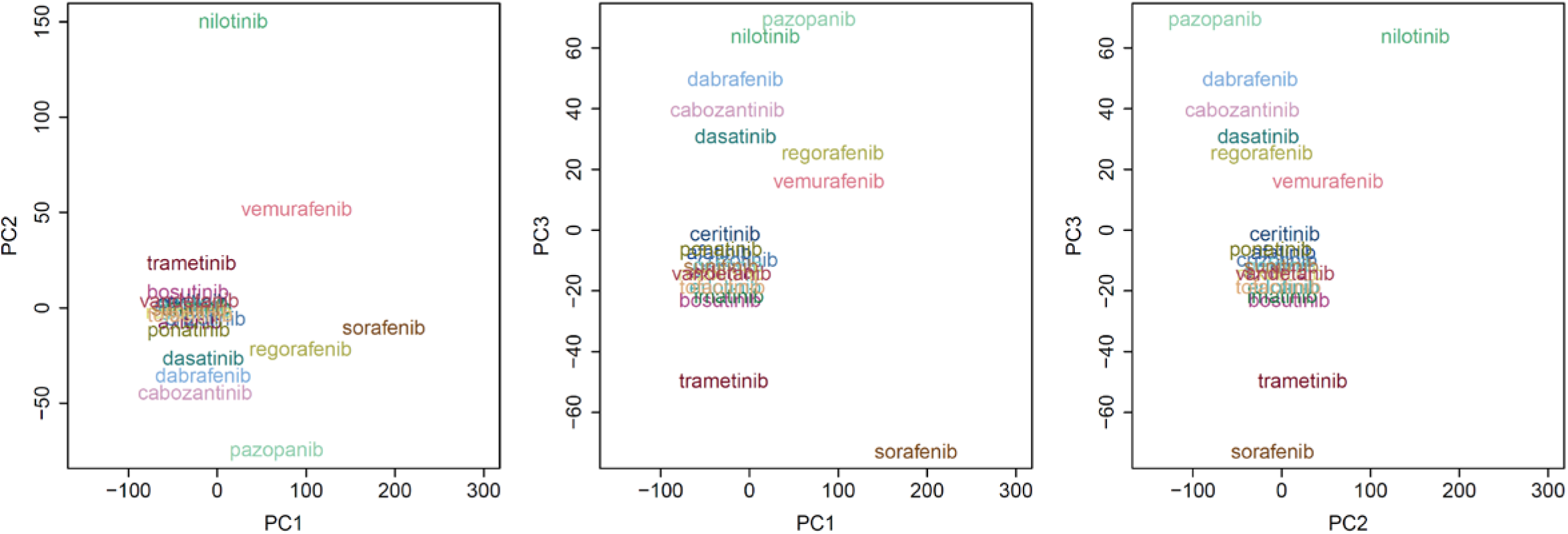
Transcriptomic profiling across KIs. For each drug, genes were ranked by mean fold-change gene expression value across replicates (>3) and cell lines, and the top 500 genes were kept. (C) First three principal components based on full mean fold-change gene expression profiles across KIs.

**Table 1.** Overview of KIs included in this analysis.

| Drug | Approval year<sup>1</sup> | Therapeutic targets | Therapeutic concentration (uM) | Cell lines |
| --- | --- | --- | --- | --- |
| Afatinib | 2013 | ErbB2, EGFR | 0.05 | 4 |
| Axitinib | 2012 | VEGFR1/VEGFR2/VEGFR3/PDGFR | 0.2 | 4 |
| Bosutinib | 2012 | Bcr-Abl, SRC | 0.1 | 2 |
| Cabozantinib | 2012 | c-Met, VEGFR2 | 2 | 3 |
| Ceritinib | 2014 | ALK | 1 | 3 |
| Crizotinib | 2011 | ALK, HGFR | 0.25 | 4 |
| Dabrafenib | 2013 | BRAF | 2.5 | 4 |
| Dasatinib | 2006 | ABL, ARG, KIT, PDGFR$\alpha/\beta$, SRC | 0.1 | 3 |
| Erlotinib | 2004 | ErbB1 | 3 | 3 |
| Gefitinib | 2003 | ErbB1 | 1 | 4 |
| Imatinib | 2001 | Bcr-Abl | 5 | 4 |
| Nilotinib | 2007 | Bcr-Abl | 3 | 4 |
| Pazopanib | 2009 | VEGFR2, PDGFR$\alpha/\beta$, KIT | 10 | 3 |
| Ponatinib | 2012 | Bcr-Abl, BEGFR, PDGFR, FGFR, EPH, SRC, c-KIT, RET, TIE2, FLT3 | 0.1 | 3 |
| Regorafenib | 2012 | RET, VEGFR, PDGFR | 1 | 2 |
| Ruxolitinib | 2011 | JAK | 1 | 3 |
| Sorafenib | 2005 | B-RAF, VEGFRs, PDGFR$\alpha/\beta$, FLT3, | 0.5 | 2 |
| Sunitinib | 2006 | VEGFR, PDGFR, CSF1R, FLT3, KIT | 1 | 3 |
| Trametinib | 2013 | MEK1, MEK2 | 0.1 | 3 |
| Tofacitinib | 2012 | JAK | 1 | 3 |
| Vandetanib | 2011 | RET, VEGFR, EGFR | 0.33 | 2 |
| Vemurafenib | 2011 | BRAF | 2 | 3 |

<sup>1</sup>US Approval date, first indication; See Table S3 for literature references to therapeutic concentrations.

### Clinical risk profiles derived from FAERS show differences in risk profiles between cardiotoxicity subtypes

To obtain estimates of clinical risk of KI-associated CT, we analyzed data from the publicly accessible FDA Adverse Event Reporting System (FAERS) (Figure 2A). We distinguished between four relevant subtypes of CT that can be identified from the FAERS database: hypertrophic cardiomyopathy (HCM), dilated cardiomyopathy (DCM), ventricular dysfunction and heart failure (VDHF), and cardiotoxicity undefined (CTU). Subsequently, reporting odds ratios (RORs) and Z-scores were derived based on the relative frequencies of CT, for each KI present in the transcriptomics dataset (Figure 2B). For several KIs, there are observable differences in the risk for different subtypes of CT. The Z-scores were used as single metric to quantify CT risk, as it includes the confidence interval for the ROR. Here, the estimates for the ROR and associated Z-score indicate relative ranking of KI-associated toxicity, and not absolute risks of CT. Finally, using multivariate logistic regression analyses we estimated the effect of age and sex at 1.02 year^-1^ (2.6 CV%) and 1.47 (1.3 CV%), respectively. The low inter-KI variation of <2.6 CV% suggest that age and sex have negligible impact on individual KI-associated CT risk estimates.

**Figure 2.**
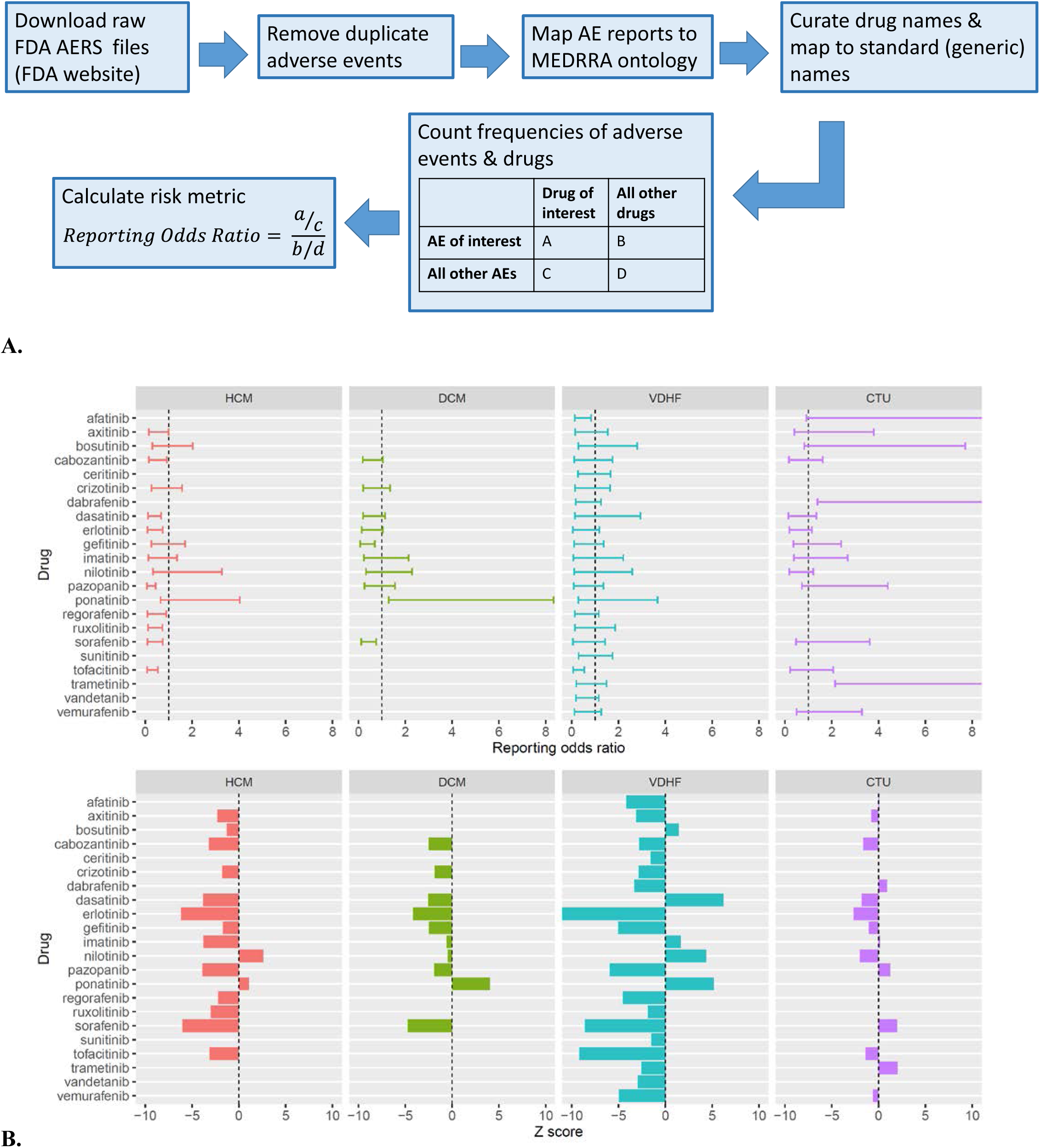
(A) Overview of data mining procedure to obtain FAERS based CT risk scores. (B). FDA Adverse Event Reporting System (FAERS) derived reporting odds ratios (top) and associated Z scores (bottom) for four different subtypes of cardiotoxicity: hypertrophic cardiomyopathy (HCM), dilated cardiomyopathy (DCM), ventricular dysfunction and heart failure (VDHF), and cardiotoxicity undefined (CTU), for different kinase inhibitors (KIs). A reporting odds ratio >1, or, a Z-score > 0 indicates above average risk for CT.

### Distinct and predictive signatures for 4 types of cardiotoxicity

Subsequently, we aimed to associate our KI-wide mean fold change gene expression profiles with the KI-specific clinical risk scores of four subtypes of CT (i.e. HCM, DCM, CTU, VDHF). Given the limited similarity between top ranking gene expression profiles across KIs (Figure 1A-C), the entirety of the gene expression profile across KIs were considered as potential predictors for KI-associated CT risk. As such, KI-specific expression profiles of 9,281 genes were available as potential predictors for 22 KI-specific CT risk scores. To identify genes most strongly associated with CT risk we used an elastic net penalized regression approach, which is suited to identify predictors whilst limiting the risk of overfitting^20^. Separate models were fit for all four derived subtypes of CT. A two-stage regression analysis was performed (Figure 3). In the first stage, we constructed bootstrap datasets obtained after random resampling of KI risk and associated gene expression profiles, which were subsequently fit as elastic net models. This first step was performed to identify gene-based predictors that could consistently predict CT risk and contributed significantly to the prediction of this risk. The bootstrap analysis resulted in stable selection of potential predictors after 250 bootstrap samples (Figure S1). Predictors to be included in the final elastic net regression model (**Supplemental material 4**) were selected based on their minimal root mean squared prediction error (RMSE, Figure S2–3) after cross validation. Since we had only a limited number of KIs available, we explicitly chose to not keep apart any data for a full external validation. Instead, we evaluated model predictions using repeated random 3-fold cross validation for each of the four subtypes of CT, which resulted in high R^2^ prediction metrics (>0.87) (Figure 4A). The selected gene expression-based predictors in the final elastic net model consisted of 42, 44, 44 and 85 genes for HCM, CTU, VDHF and DCM, respectively (Table S2, Figure 4B). These sets of genes define our clinically-weighted transcriptomic signatures for cardiotoxicity; their associated gene expression profiles are shown in Figure 4C. The importance of each of the selected genes for the prediction of CT risk was not equal (Figure 4B). CT subtype DCM required 85 genes for optimal predictive performance, whereas for all other subtypes <44 genes were sufficient. Potentially, this is because not enough KIs with a diverse CT profile were available for deriving an informative gene signature. When assessing the similarity between each of the 4 clinically weighted gene expression signatures, the largest overlap was present between DCM and HCM (4 genes), VDHF and DCM (4 genes), and VDHF and HCM (2 genes) (Figure 4D). The genes FASTKD2 (FAST kinase domain containing protein 2) and MED 19 (Mediator Of RNA Polymerase II Transcription, Subunit 19) were present in the signatures for HCM, VDHF and DCM. FASTKD2 plays a role in energy balance regulation in mitochondria under stress^21^. This could indeed be of relevance for cardiotoxicity, because mitochondrial dysfunction has been associated with cardiotoxicity^13,22^, and this finding may warrant further detailed analysis.

**Figure 3.**
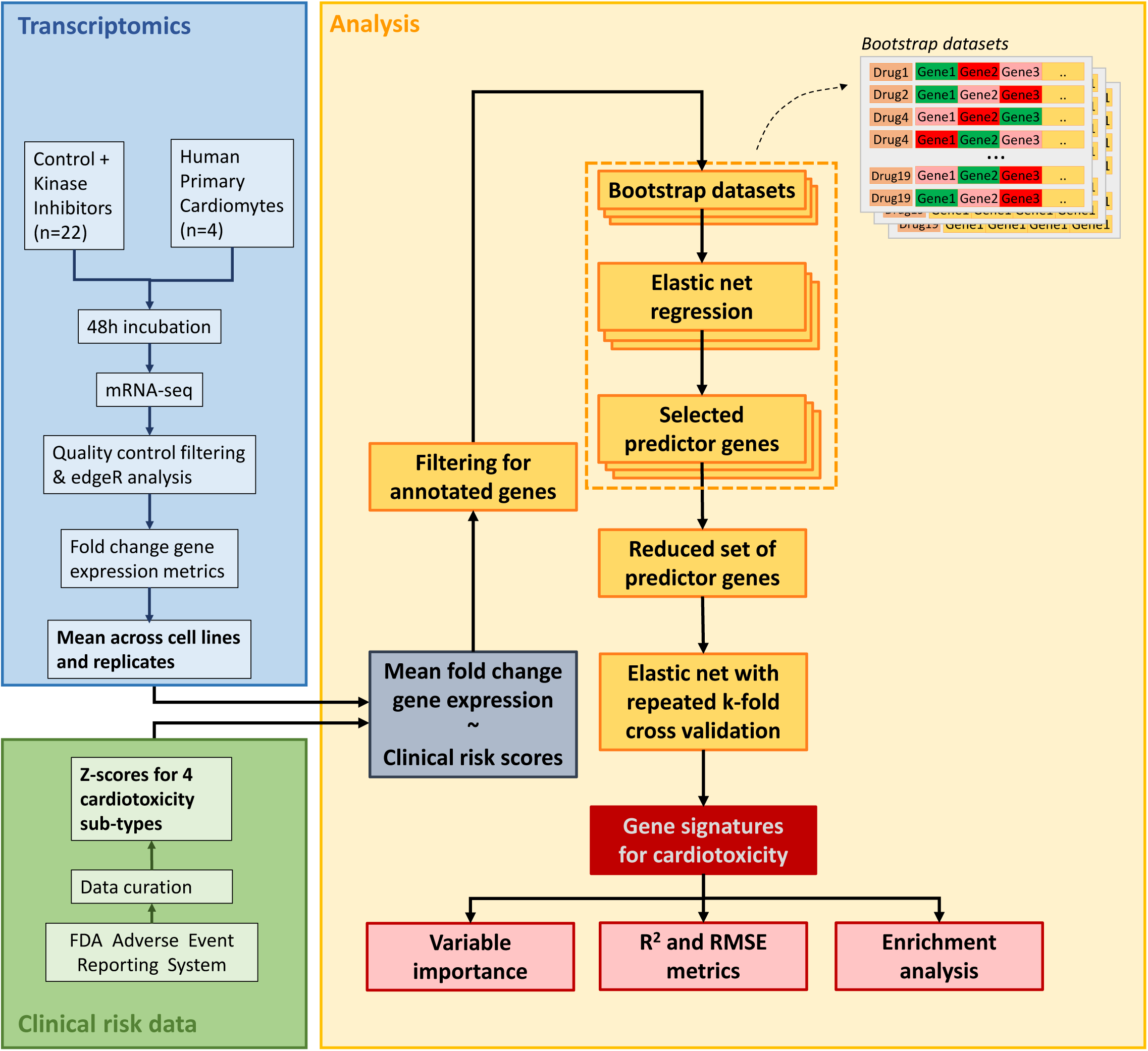
Schematic overview of the methods used for data generation and analysis.

**Figure 4A.**
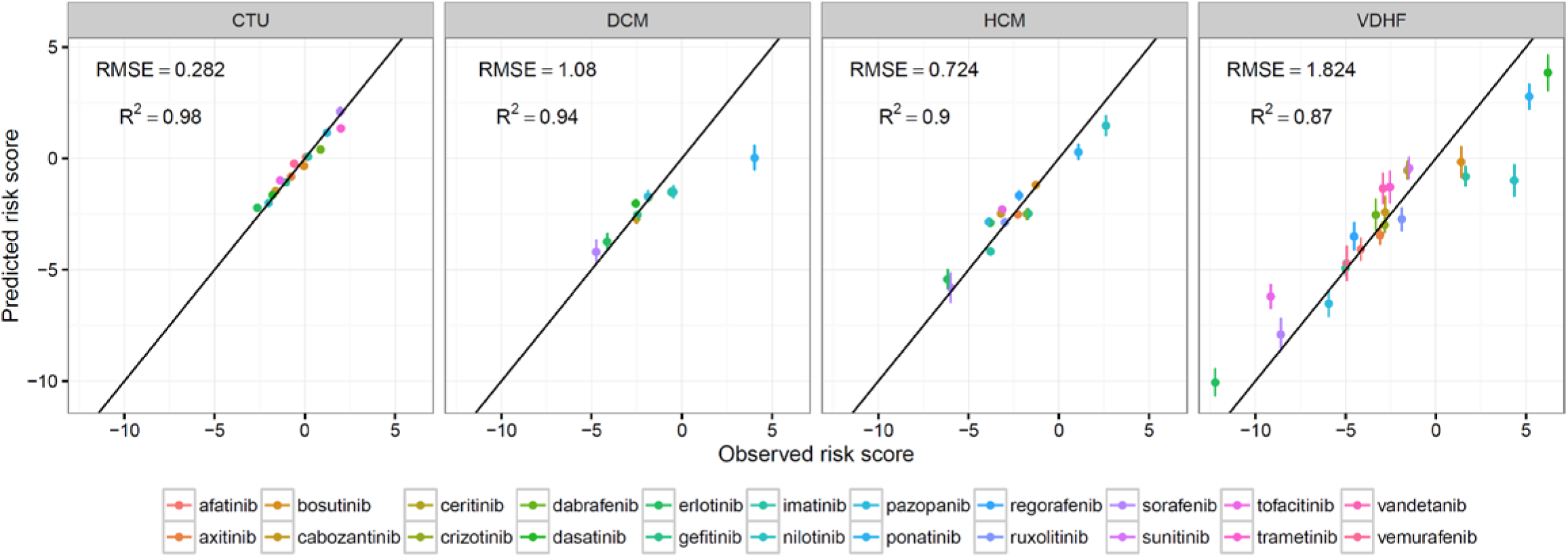
Regression analysis and signatures derived from clinical risk scores and transcriptomics profiling. (A) 3-fold cross validation based mean and standard deviation of predicted and observed clinical Z-score metrics, and their associated mean R^2^ and RMSE metrics for four subtypes of cardiotoxicity.

**Figure 4B.**
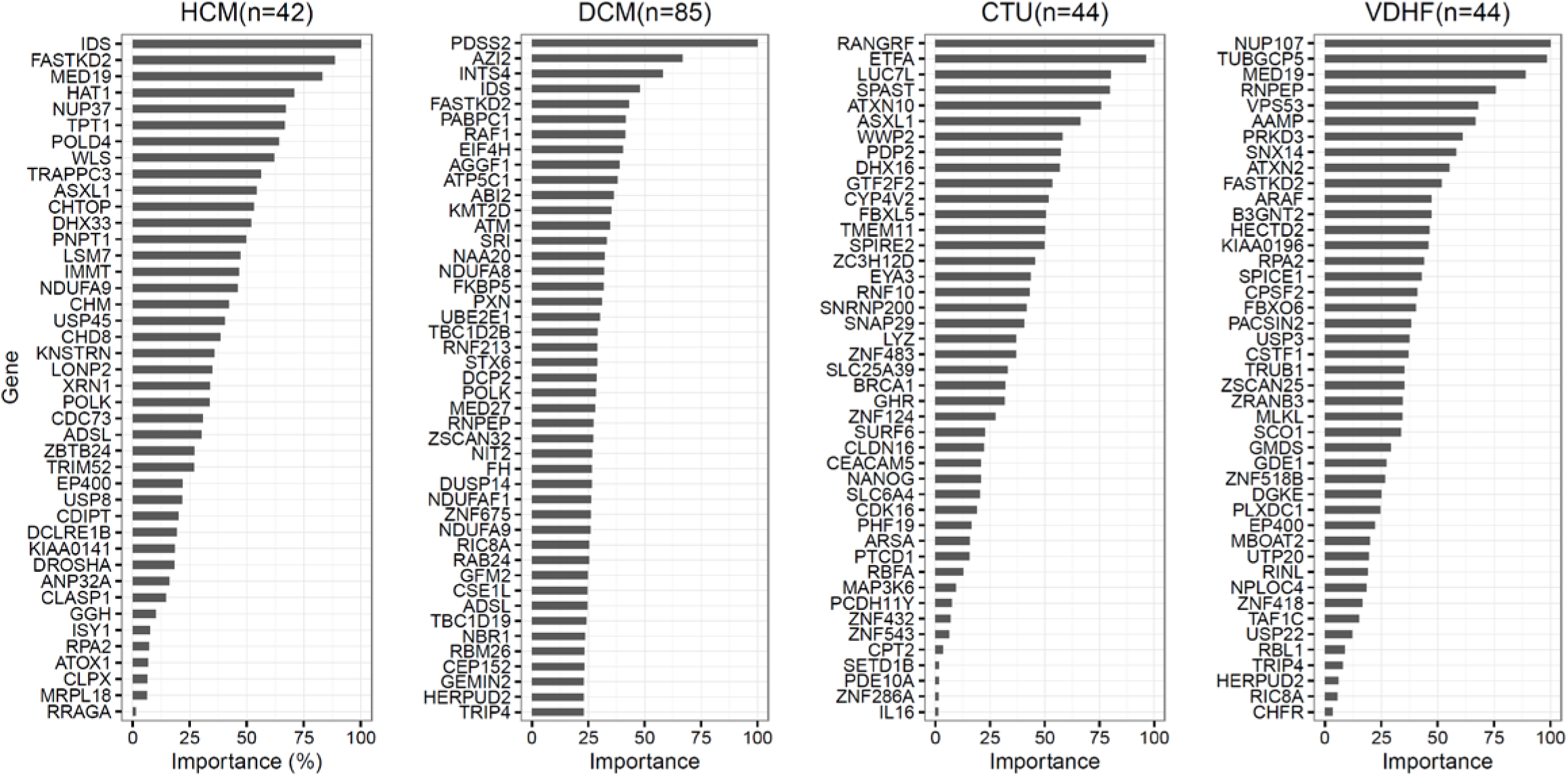
Regression analysis and signatures derived from clinical risk scores and transcriptomics profiling. (A) 3-fold cross validation based mean and standard deviation of predicted and observed clinical Z-score metrics, and their associated mean R2 and RMSE metrics, for four subtypes of cardiotoxicity. (B) Variable importance ranking of final genes signatures, with number of total genes in signature.

**Figure 4C.**
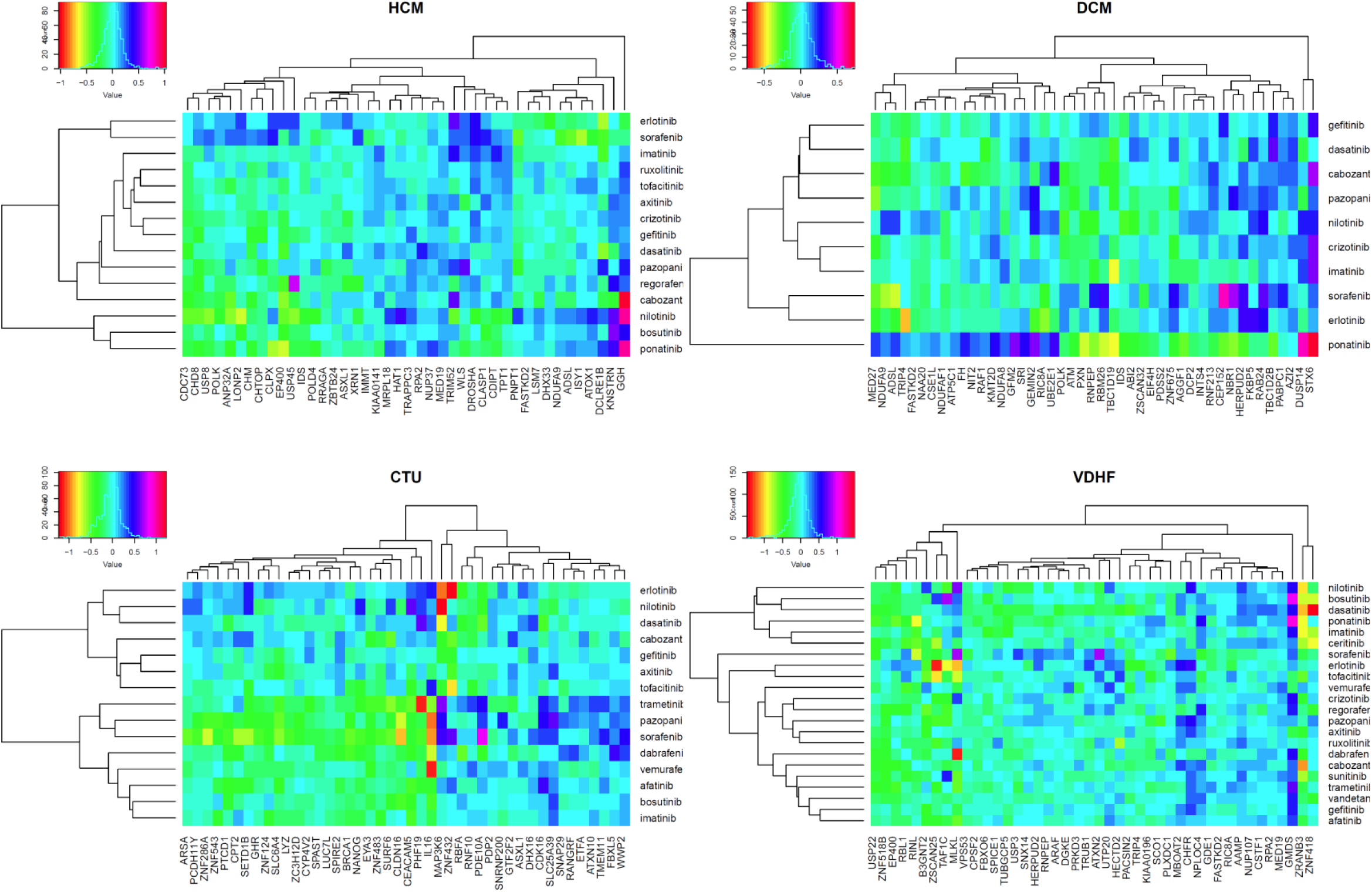
Clustered heatmaps of selected predictors and associated mean fold change gene expression values for different KIs. In plots B and C, for DCM, the lowest ranking genes were not included. These were: ZKSCAN7, ANAPC4, COPS4, SLC25A3, DAB2, PQLC2, ING5, MRPL18, NUP37, CLPX, MRPL51, RBFA, ZDHHC20, ERBB2IP, MBOAT2, MALSU1, PLK2, GMDS, NADSYN1, WLS, ATG12, ENY2, ZNF180, NR2C2, MED19, FAM175B, DLD, ZNF702P, SUCLG1, TIA1, ZNF28, PML, CBLB, ZMYM4, ZSWIM7, KMT2E, NDUFA4, DST, ATG7, PCDHB7.

**Figure 4D.**
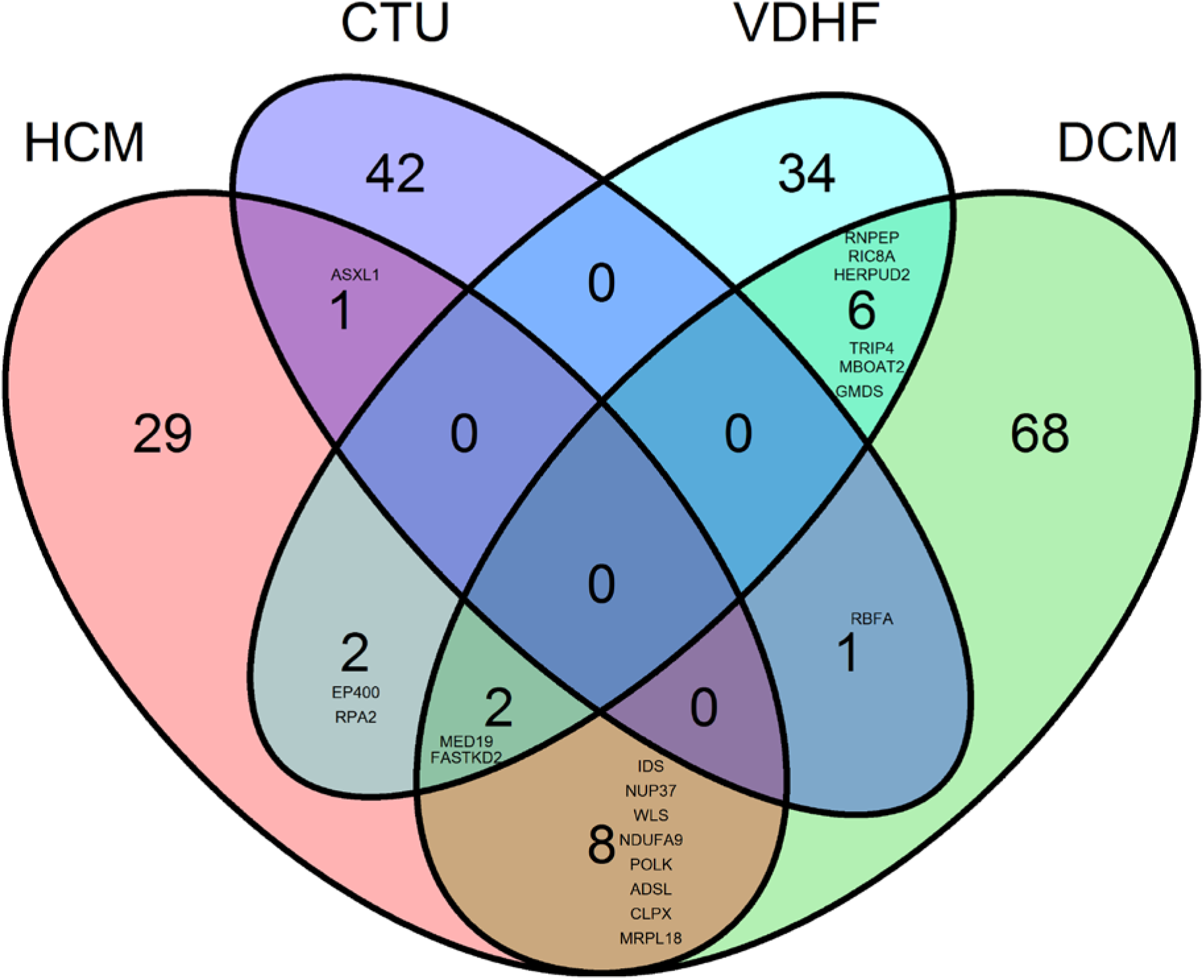
Venn diagrams of gene selected in the final gene signature. Overlapping genes have been indicated.

### Signatures for the 4 types of cardiotoxicity are enriched for different biological processes

In order to identify the potential functional consequences of our clinically-weighted transcriptomic signatures, we performed enrichment analyses based on the KEGG database, estimating over-representation of signature genes using Fisher’s exact test (Figure 5). Biological processes that appeared to be affected for all four types of CT included metabolic pathways, and common subcellular processes such as ribosome function, endocytosis, proteolysis, and splicing. However, similar to the differential clinical risks for the different types of cardiotoxicity, we observed distinct patterns in enriched subcellular processes for specific CT subtypes. Such differences include regulation of the actin cytoskeleton and oxidative phosphorylation for DCM, protein processing in the endoplasmic reticulum for HCM, VDHF), and apoptosis (DCM) amongst others (Figure 5).

**Figure 5.**
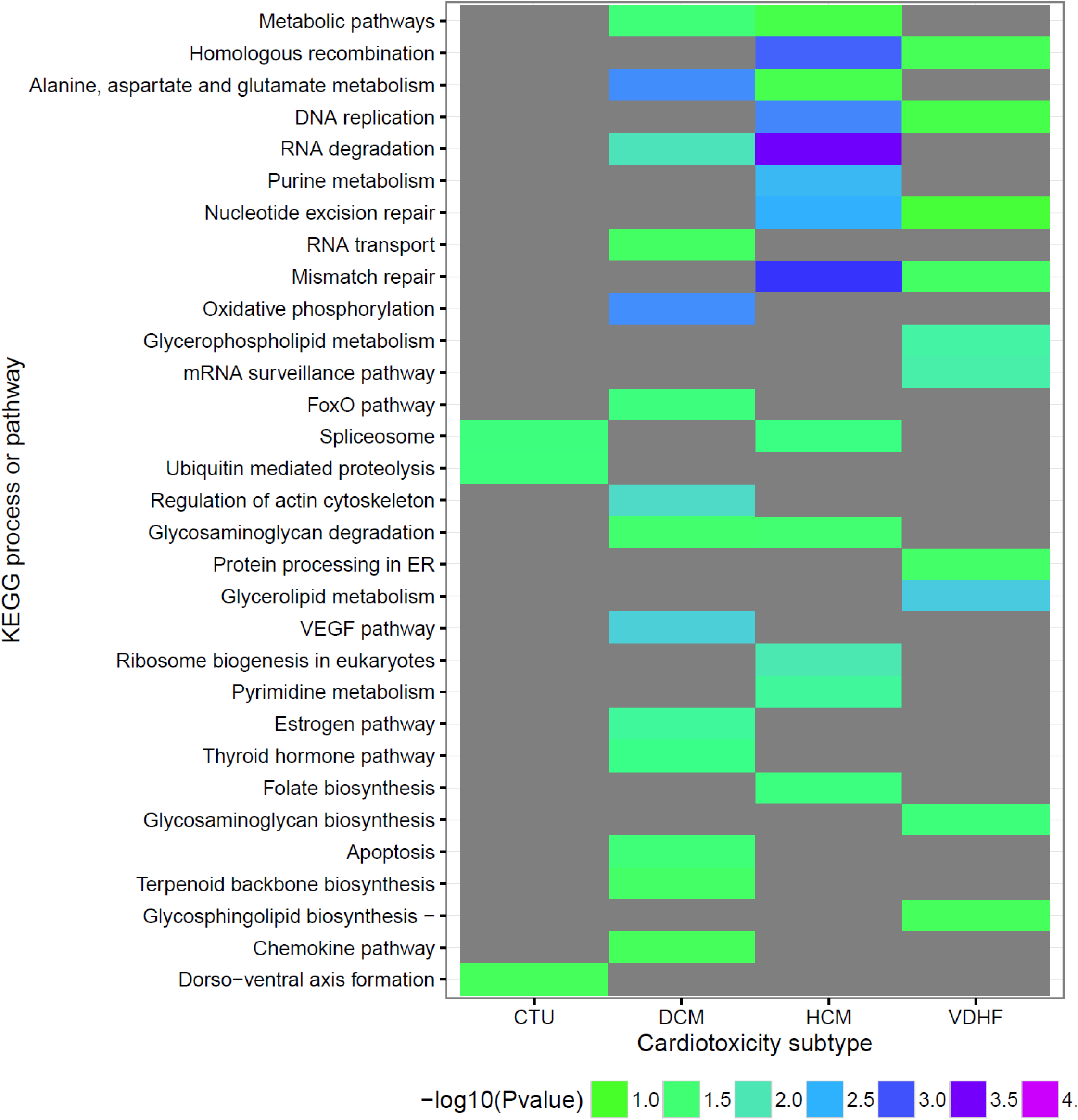
Enrichment analysis based on the KEGG database of clinically weighted gene expression signatures for cardiotoxicity subtypes ranked by their combined significance.

## DISCUSSION

The occurrence of drug treatment associated cardiotoxicity leading to decreased cardiac contractility lags behind the therapeutic effects of the drugs and may only be observed in a subset of the patients using the drug. These two factors raise the question of whether it would be possible to obtain early cell based indicators of potential for drug toxicity. This study was designed to address this question. The approach we have used is to determine we can associate drug treatment induced gene expression patterns and the clinical risk for the adverse events of interest. We estimated clinical risk from the FAERS database. Our use of FAERS is arguably a relevant and unbiased approach for the quantification of CT risk, because this data reflects unselected, real-life patients cohorts and may some of the potential bias associated with cohort selection associated with tightly controlled clinical trials. It is possible that risk metrics derived from controlled clinical trials can underestimate adverse event risks due to selective inclusion criteria of patients to demonstrate therapeutic efficacy, cohort size and duration of the trial. Nevertheless there are limitations to the FAERS database as well that we have discussed and addressed in previous work^18^, and similarly address in this study. In particular, we confirmed that demographics such as age and sex were not variable across different KIs. Moreover, as the KIs studied are cancer drugs, some similarity between populations is likely.

This project associated KI-associated transcriptomic response profiles generated from cultured human primary cardiomyocytes with clinical risk scores for different subtypes of CT in order to obtain a reduced set of genes that may predict the relative risk for KI-associated CT. The clinically weighted transcriptomic signatures consisted of <44 genes for HCM, CTU, and VDHF, whilst the signature for DCM consisted of 85 genes. The signatures showed good prediction of CT risk based on the cross validation analysis (Figure 4A). Using the clinically weighted signatures and the association regression coefficients identified in the elastic net model (**Supplemental material 4**), the relative risk for CT can be predicted. Here, the risk metric does not reflect the absolute risk for developing CT. Rather it reflects the relative risk for a subset of patients for which drug-associated adverse events were reported. As such, the risks predicted by our signatures and associated regression model can be used in drug development to rank the risk of potentially novel KIs with respect to the current ranking of existing KIs with better characterized clinical risks for CT. The issue of whether or not a KI can be considered as cardiotoxic has some underlying complexity. Some KIs may result in cardiotoxicity, but the severity and duration of CT may be different between patients and between different KIs. Therefore, we decided to develop a continuous risk metric in preference to a binary classification of drugs as CT or not. Secondly, it is unclear if all KIs induce CT through similar mechanisms and to what extent ultimate clinical pathologies are similar. Indeed, the FAERS database allows us to distinguish between different types of CT, rather than lump all potential types of CT together (Figure 2B). Nonetheless, the clinical classification of KI associated CT is an area that needs better resolution, and it is likely as EHR records become increasingly available that the data will become more detailed with respect to pathophysiology. Such improved clinical data can greatly enhance the value of analyses such as ours and aid in the drug development processes by contributing to early go/no-go decisions.

At the cellular level, commonly enriched subcellular processes associated with the clinically-weighted transcriptomic signatures included various metabolic processes, pathways related to oxidative phosphorylation, endoplasmic reticulum (ER) effects, and apoptosis. In addition, a number of subcellular processes related to transcription (ribosome, spliceosome), protein degradation and endocytosis were affected. Other recent studies into CT of anthracyclines and KIs agree with our findings that changes in metabolic processes^23,24^, ER effects^14^, oxidative phosphorylation^13,25^ AMPK signaling^26^, cell adhesion^27^, cell survival pathways^12^, and the FoxO-related transcription^28^ are likely to be involved. It is possible that gene expression profiles associated with specific subcellular processes could be co-expressed and hence highly correlated. However, due to the nature of the elastic net regression algorithm used, highly correlated genes are unlikely to be selected simultaneously, limiting over representation of the identification of associated subcellular pathways.

Besides variation in drug exposure, pharmacodynamic inter-individual variation in severity of CT is likely. The current group of four cell lines are not of sufficient size to realistically capture such human variability. Therefore, in our analysis, we used mean fold change gene expression profiles across multiple cell lines. The resulting averaged gene expression profiles thus reflect relatively consistent changes in gene expression across cell lines, i. e. changes in gene expression that are less likely to be highly variable across individuals, yet may result in consistent predictors across the population. Given that the FAERS CT risk scores also reflect a population-level CT risk, the use of these mean values in fold-change gene expression values is a reasonable starting point for our analyses.

The experimental underpinning of the transcriptomic profiles generated in this study make them likely to be of value in selecting drug candidates for human use. Our analysis is based on primary human derived cardiomyocytes. Although these cell lines do have phenotypic limitations due to dedifferentiation^29^, the signatures obtained from the cells could be relevant for prediction of clinical drug effects. These cell lines may be reflective of specific human cardiac pharmacology^30,31^, even though further characterization and standardization is still needed. Our analyses used drug exposures at estimated therapeutic concentrations of the individual KIs, rather than using the same concentrations for all KIs. We used the reported maximum concentrations as reported in literature (Table S3). Here, we did not correct for protein binding, but, we consider that given the typical protein binding of >95% of these KIs, the concentration used may reflect patients with over-exposure to the KI. As such, the concentrations used are relevant to assess transcriptomic changes that are likely related to early changes in subcellular processes associated with the adverse event of interest.

We anticipate that clinically weighted transcriptomic signatures such as developed in this study may be of relevance to guide safety assessment in early drug development. Unlike the relatively well-established assessment of electrophysiological safety issues such as QT prolongation, the assessment of non-QT type of cardiotoxicity associated with KIs^12^ and other novel drugs^32^, lack reliable biomarkers. Transcriptomic signatures could help fill this gap. Transcriptomic analyses may also guide the discovery of subcellular pathway based biomarkers that can be measured in both preclinical model systems as well as in patients in surrogate cells. Data on biomarker levels could then be included in PK-PD models to personalize dose regimens that limit risk of CT^33–35^, before the appearance of imaging based indicators such as the left ventricular ejection fraction. The current study represents a starting point and many more studies both experimental and computational will be needed to utilize transcriptomic studies in drug development and clinical practice.

## METHODS

### Experiments to identify differentially expressed genes in human cardiomyocytes treated with KIs

Adult human cardiomyoctes were purchased from PromoCell and grown in culture per manufacturer’s instructions. Four different cell lines representing different human subjects were studied. Cells were treated with a single dose at the estimated therapeutic concentration used in humans for 48 hrs (Table S3). After drug treatment, the cells were lysed, RNA was collected using TRIzol, and gene expression profiles were measured using the 3’ digital gene expression method^36,37^. Details of the experimental protocols have been described in another study^38^ and step-by-step standard operating procedures for the various experiments are available on www.dtoxs.org.

### Sequence alignment and processing of gene expression data

The raw sequences were demultiplexed. Combined standard RNA-seq sequencing files were aligned to the reference human genome provided by the Broad Institute^36^, using the Burrows-Wheeler Alignment (BWA)^39^ software. Details of these computational procedures are described elsewhere^38^, and step-by-step protocols are available on ww.dtoxs.org. The resulting alignment files were parsed to identify the fragments with acceptable alignment quality, to remove duplicate fragments, and to assign accepted fragments to corresponding genes. The resulting read-count (i.e., transcript count) table was then subjected to correlation analysis at each treatment condition, to identify and remove outlier samples, determined by predefined thresholds. The gene read-count tables were then subjected to differential gene expression analysis using the R package EdgeR^40^. The resulting normalized and log-transformed fold-change gene expression values for each sample were deposited for public access to the DtoxS repository (www.dtoxs.org).

### Processing and exploratory analysis or gene expression data

The mean log transformed gene expression fold-change value was calculated across all cell lines for each individual KI. The resulting matrix of gene fold changes values by KIs was used for the regression analysis. To obtain insight in the general patterns present in this KI-perturbed transcriptomics dataset, we generated rankings of the top 500 genes for each drug, by their absolute fold change value. For each of these KI-associated rankings we determined the frequency of these changes being also present in the ranking of other drugs, e.g. the similarity in top ranking gene expression. This was visualized using the Jaccard index, and by plotting the most highly drug-connected genes against the associated drugs. Finally we performed on the mean fold-change values for each drug a principal component analysis for the first 3 principal components to further assess similarity between drugs in their gene expression values.

### Calculation of clinical risk scores

The FDA Adverse Event Reporting Systems (FAERS) raw data files were downloaded from the FDA website (all report files up to 2015 Q3). Adverse events in the raw data files were mapped to the MEDDRA dictionary^41^. Subsequently we defined four types of CT, that were related to either hypertrophic cardiomyopathy, dilated cardiomyopathy, ventricular dysfunction and heart failure, and undefined cardiotoxicity. Electrophysiological forms of CT were excluded. The selection of the selected CT subtypes is provided in Table S1. Since drug names in FAERS are of low quality, i.e. they contain many spelling mistakes, mixed brand names, or combinations of drugs associated with a single adverse event, we performed extensive curation on the drug names converting the drug name records to generic drug names. This was done using dictionaries from the FDA and DrugBank. We also applied a conservative fuzzy match algorithm to detect drug names with spelling errors. Through iterative manual checking and automated string matching using R we derived a curated database of drug names associated with adverse events. Subsequently, the frequency of CT events in the 4 predefined subgroups, other non-CT adverse events were computed for individual KIs included in our experiments, and for all other drugs. Based on this we calculated a reported odds ratio (ROR) as follows, ROR = (a/b)/(c/d), where a, b, c, and d are the number of individuals with a drug-associated adverse event, where a is the number of individuals who received drug X, and had the adverse event of interest, b is the number of individuals who received drug X and did not have the adverse event of interest; c is the number of individuals who did not receive drug X but some other drug, and had the adverse event of interest; and d is the number of individuals who did not receive drug X and did not have the adverse event of interest. Subsequently, the standard error of the log transformed odds ratio (SElogROR) was computed as follows: SE=sqrt(1/a+1/b+1/c+1/d). The Z-score was then computed as Z=log(ROR)/SElogROR. To assess the impact of age and sex as confounders variable for different KIs, we performed multivariate logistic regression analyses for each of the individual KIs in combination with factors estimated for age and sex. For this we used the standard generalized linear model available in R.

### Elastic net regression analysis

The FAERS-derived risk (Z-scores) for each of the four types of CT was regressed against the KI associated vectors of mean fold change values across the four cell lines. We chose to use Z-scores as metric reflecting CT-risk, as this metric takes into consideration the uncertainty of the reporting odds ratio. Prior to regression we filtered the gene lists by their presence in either the Gene Ontology Biological Processes, KEGG, Reactome, or WikiPathways databases, in order to exclude genes for which no biological function has been established. Subsequently we generated 250 bootstrap datasets with replacements. We did separate bootstrapping for each KI. Each of these bootstrap datasets was fit using an elastic net regression model (R package glmnet). The selected genes that were selected as predictors (i.e. non-zero regression coefficient) and the scaled values of the gene-associated coefficients were saved for each bootstrap dataset. Across all bootstrap datasets the relative frequency of selection of gene-based predictors, and the mean scaled coefficient value of these coefficients was computed. We then calculated the product of the mean frequency and scaled coefficient value, rank predictors by their importance with respect to robustness (selection frequency) and their importance. Different upper percentiles of these rankings were regressed against each CT subtype and evaluated using 3-fold cross validation. The selection percentile resulting in optimal prediction errors (RMSE) was then used to select a subset of gene based predictors, and to select the model that generated the final gene expression signatures for each CT subtype. The selected predictor genes were then visualized ranked by their relative importance, and by their mean fold change values as clustered heatmaps.

### Enrichment analyses

Enrichment analysis was performed based on a one-tailed Fisher’s exact test using R, in order to identify enrichment of specific genes in predefined gene lists, as present in the KEGG database. We performed enrichment of the selected gene-based predictors for the four types of CT, as present in the final signatures. Diseases were excluded from the KEGG list of processes (e.g. diabetes, depression, cancer), in order to only evaluate general biological processes or pathways.

## ACKNOWLEDGEMENTS

This project was supported in part by the NIH LINCS center grant (U54HG008098) and the Systems Biology Center grant P50GM071558. JGCvH received funding from the European Union MSCA program (Project ID 661588).

## CONFLICT OF INTEREST

The authors declare no conflict of interest.

## SUPPLEMENTAL MATERIALS. TABLES

**Table S1.**
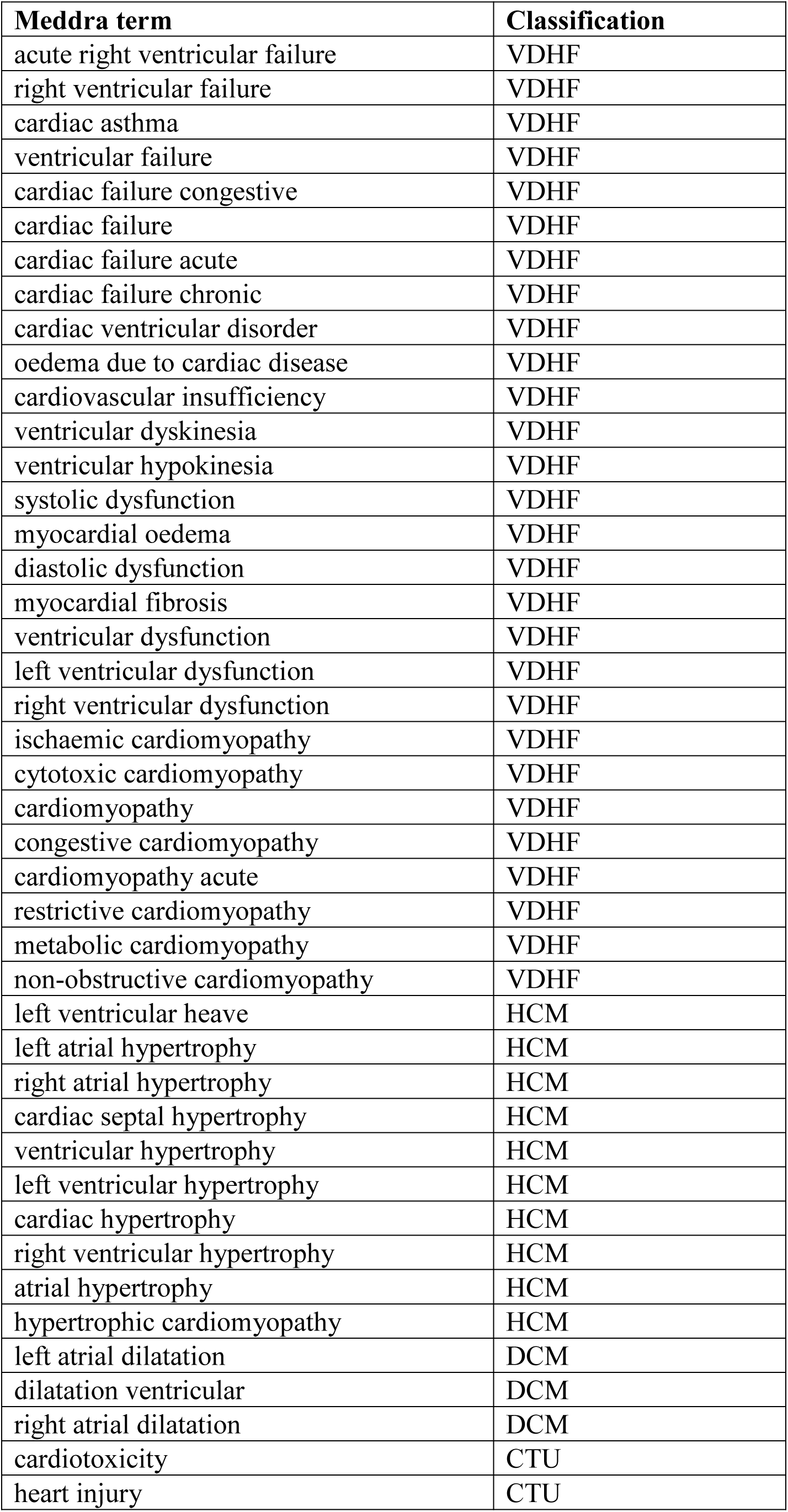
Definition of MEDDRA subgroups for different cardiotoxicity subtypes used.

| **Meddra term** | **Classification** |
| --- | --- |
| acute right ventricular failure | VDHF |
| right ventricular failure | VDHF |
| cardiac asthma | VDHF |
| ventricular failure | VDHF |
| cardiac failure congestive | VDHF |
| cardiac failure | VDHF |
| cardiac failure acute | VDHF |
| cardiac failure chronic | VDHF |
| cardiac ventricular disorder | VDHF |
| oedema due to cardiac disease | VDHF |
| cardiovascular insufficiency | VDHF |
| ventricular dyskinesia | VDHF |
| ventricular hypokinesia | VDHF |
| systolic dysfunction | VDHF |
| myocardial oedema | VDHF |
| diastolic dysfunction | VDHF |
| myocardial fibrosis | VDHF |
| ventricular dysfunction | VDHF |
| left ventricular dysfunction | VDHF |
| right ventricular dysfunction | VDHF |
| ischaemic cardiomyopathy | VDHF |
| cytotoxic cardiomyopathy | VDHF |
| cardiomyopathy | VDHF |
| congestive cardiomyopathy | VDHF |
| cardiomyopathy acute | VDHF |
| restrictive cardiomyopathy | VDHF |
| metabolic cardiomyopathy | VDHF |
| non-obstructive cardiomyopathy | VDHF |
| left ventricular heave | HCM |
| left atrial hypertrophy | HCM |
| right atrial hypertrophy | HCM |
| cardiac septal hypertrophy | HCM |
| ventricular hypertrophy | HCM |
| left ventricular hypertrophy | HCM |
| cardiac hypertrophy | HCM |
| right ventricular hypertrophy | HCM |
| atrial hypertrophy | HCM |
| hypertrophic cardiomyopathy | HCM |
| left atrial dilatation | DCM |
| dilatation ventricular | DCM |
| right atrial dilatation | DCM |
| cardiotoxicity | CTU |
| heart injury | CTU |

**Table S2.** Final gene signatures for four subtypes of CT.

| DCM | HCM | CTU | VDHF |
| --- | --- | --- | --- |
| PDSS2<br>AZI2<br>INTS4<br>IDS<br>FASTKD2<br>PABPC1<br>RAF1<br>EIF4H<br>AGGF1<br>ATP5C1<br>ABI2<br>KMT2D<br>ATM<br>SRI<br>NAA20<br>NDUFA8<br>FKBP5<br>PXN<br>UBE2E1<br>TBC1D2B<br>RNF213<br>STX6<br>DCP2<br>POLK<br>MED27<br>RNPEP<br>ZSCAN32<br>NIT2<br>FH<br>DUSP14<br>NDUFAF1<br>ZNF675<br>NDUFA9<br>RIC8A<br>RAB24<br>GFM2<br>CSE1L<br>ADSL<br>TBC1D19<br>NBR1<br>RBM26<br>CEP152<br>GEMIN2<br>HERPUD2<br>TRIP4<br>ZKSCAN7<br>ANAPC4<br>COPS4<br>SLC25A3<br>DAB2<br>PQLC2<br>ING5<br>MRPL18<br>NUP37<br>CLPX<br>MRPL51<br>RBFA<br>ZDHHC20<br>ERBB2IP<br>MBOAT2<br>MALSU1<br>PLK2<br>GMDS<br>NADSYN1<br>WLS<br>ATG12<br>ENY2<br>ZNF180<br>NR2C2<br>MED19<br>FAM175B<br>DLD<br>ZNF702P<br>SUCLG1<br>TIA1<br>ZNF28<br>PML<br>CBLB<br>ZMYM4<br>ZSWIM7<br>KMT2E<br>NDUFA4<br>DST<br>ATG7<br>PCDHB7 | IDS<br>FASTKD2<br>MED19<br>HAT1<br>NUP37<br>TPT1<br>POLD4<br>WLS<br>TRAPPC3<br>ASXL1<br>CHTOP<br>DHX33<br>PNPT1<br>LSM7<br>IMMT<br>NDUFA9<br>CHM<br>USP45<br>CHD8<br>KNSTRN<br>LONP2<br>XRN1<br>POLK<br>CDC73<br>ADSL<br>ZBTB24<br>TRIM52<br>EP400<br>USP8<br>CDIPT<br>DCLRE1B<br>KIAA0141<br>DROSHA<br>ANP32A<br>CLASP1<br>GGH<br>ISY1<br>RPA2<br>ATOX1<br>CLPX<br>MRPL18<br>RRAGA | RANGRF<br>ETFA<br>LUC7L<br>SPAST<br>ATXN10<br>ASXL1<br>WWP2<br>PDP2<br>DHX16<br>GTF2F2<br>CYP4V2<br>FBXL5<br>TMEM11<br>SPIRE2<br>ZC3H12D<br>EYA3<br>RNF10<br>SNRNP200<br>SNAP29<br>LYZ<br>ZNF483<br>SLC25A39<br>BRCA1<br>GHR<br>ZNF124<br>SURF6<br>CLDN16<br>CEACAM5<br>NANOG<br>SLC6A4<br>CDK16<br>PHF19<br>ARSA<br>PTCD1<br>RBFA<br>MAP3K6<br>PCDH11Y<br>ZNF432<br>ZNF543<br>CPT2<br>SETD1B<br>PDE10A<br>ZNF286A<br>IL16 | NUP107<br>TUBGCP5<br>MED19<br>RNPEP<br>VPS53<br>AAMP<br>PRKD3<br>SNX14<br>ATXN2<br>FASTKD2<br>ARAF<br>B3GNT2<br>HECTD2<br>KIAA0196<br>RPA2<br>SPICE1<br>CPSF2<br>FBXO6<br>PACSIN2<br>USP3<br>CSTF1<br>TRUB1<br>ZSCAN25<br>ZRANB3<br>MLKL<br>SCO1<br>GMDS<br>GDE1<br>ZNF518B<br>DGKE<br>PLXDC1<br>EP400<br>MBOAT2<br>UTP20<br>RINL<br>NPLOC4<br>ZNF418<br>TAF1C<br>USP22<br>RBL1<br>TRIP4<br>HERPUD2<br>RIC8A<br>CHFR |

**Table S3.** Typical concentrations used for experiments (exposure for 48h). Concentrations were selected based on the mean maximum drug concentrations reported in clinical studies. For dose escalation studies the highest concentration was used.

| Drug | In Vitro (uM) | Reference |
| --- | --- | --- |
| Sorafenib | 0.5 | [1] |
| Sunitinib | 1 | [2][3] |
| Crizotinib | 0.25 | [4] |
| Erlotinib | 3 | [5][6] |
| Imatinib | 5 | [7] |
| Dasatinib | 0.1 | [8][9] |
| Tofacitinib | 1 | [10] |
| Gefitinib | 1 | [11] |
| Bosutinib | 0.1 | [12] |
| Vandetanib | 0.333 | [13] |
| Lapatinib | 2 | [14] |
| Nilotinib | 3 | [15][16] |
| Axitinib | 0.2 | [17][18] |
| Pazopanib | 10 | [19] |
| Ruxolitinib | 1 | [20][21] |
| Afatinib | 0.05 | [22][23] |
| Regorafenib | 1 | [24] |
| Ponatinib | 0.1 | [25] |
| Dabrafenib | 2.5 | [26][27] |
| Vemurafenib | 2 | [28] |
| Cabozantinib | 2 | [29] |
| Trametinib | 0.1 | [30] |
| Ceritinib | 1 | [31] |

## SUPPLEMENTAL MATERIAL 3. FIGURES

**Figure S1.**
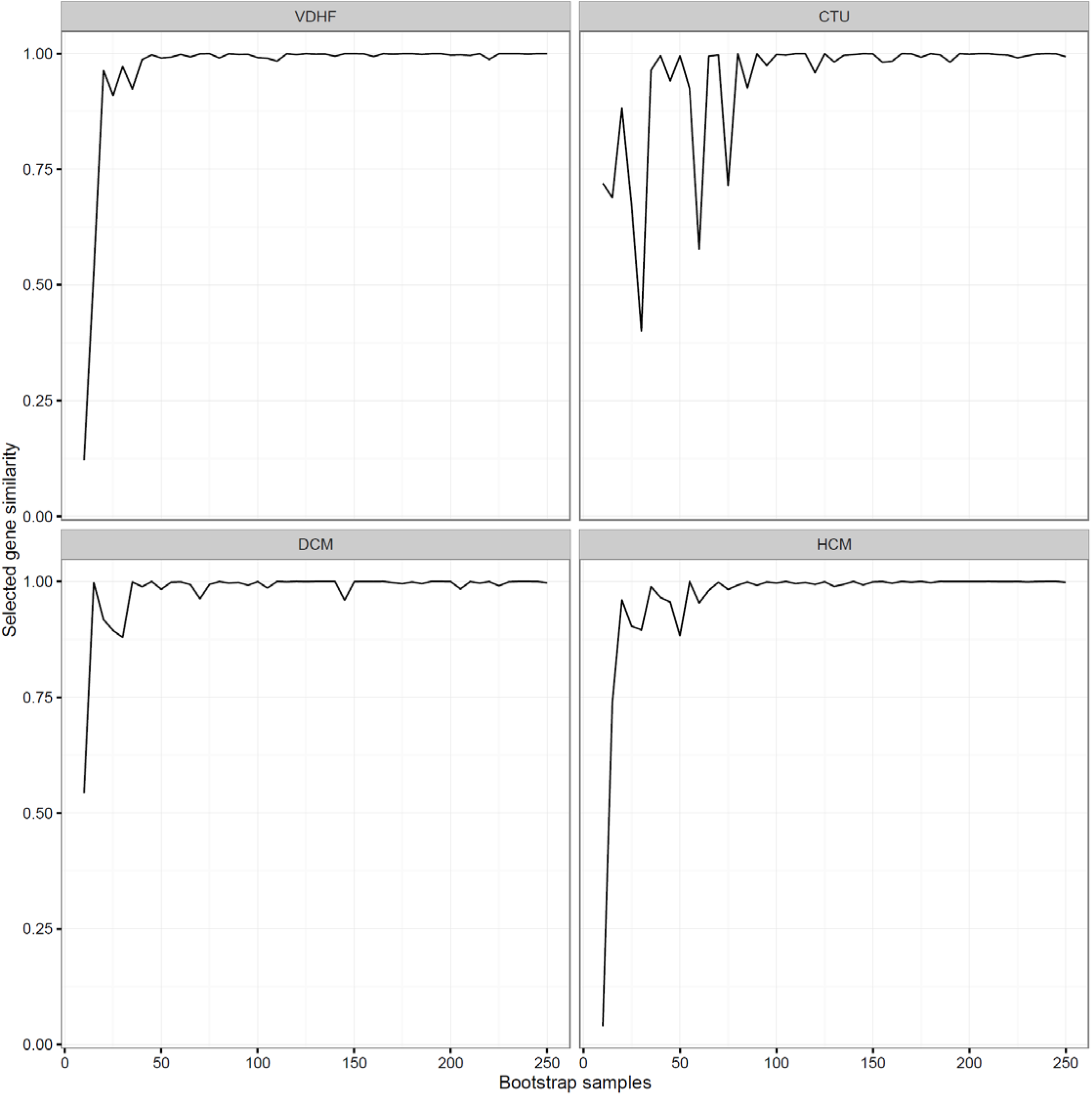
Selected gene similarity versus number of bootstrap samples for 4 sub-types of cardiotoxicity.

**Figure S2.**
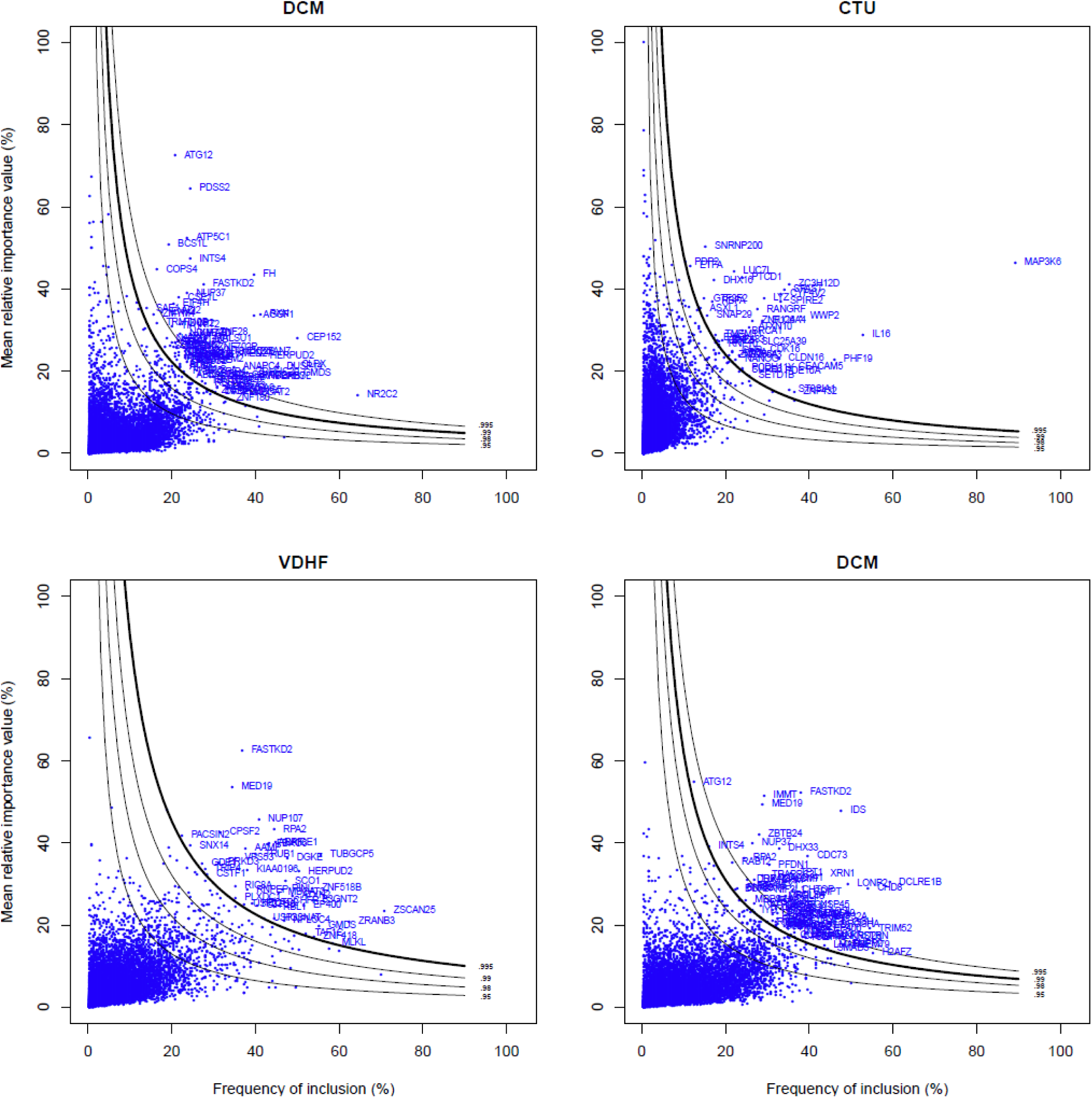
Selection of predictors from the bootstrap analysis for different CT subtypes. The blue circles indicate individual genes that were selected with their associated frequency of inclusion value and mean relative importance value, computed from the bootstrap analysis (n=250). The solid lines indicate different percentiles for the product of the frequency and the mean relative importance. The solid line with increased with indicates the optimal percentile used to fit the final gene signature regression models. The names of selected genes above the selected percentile are depicted.

**Figure S3.**
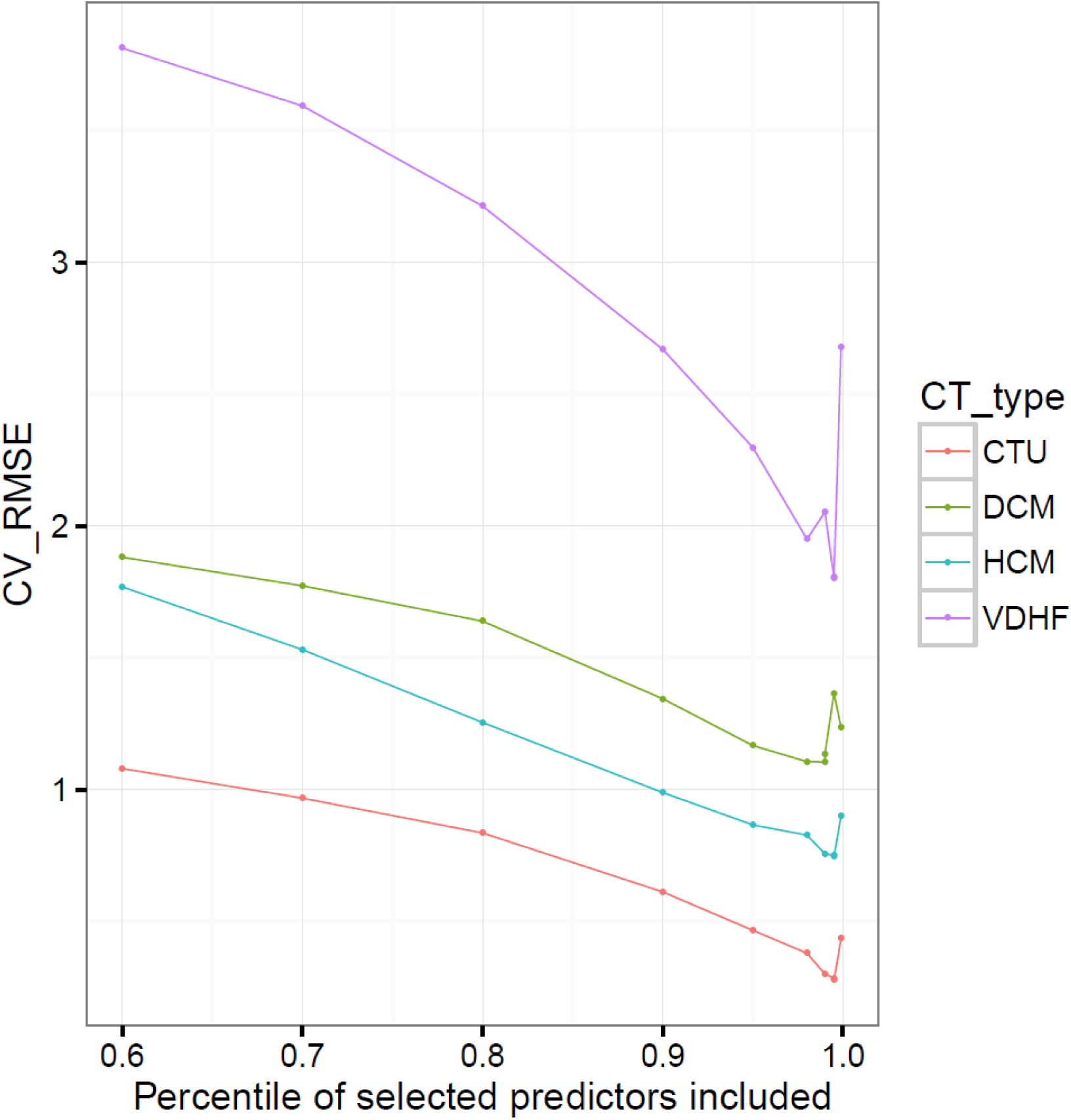
Selection of optimal threshold percentile for inclusion of genes in the final gene signature regression analysis for different subtypes of cardiotoxicity, based on the cross-validation RMSE obtained. The percentile was selected for the minimum RMSE.

